# A framework for identifying fertility gene targets for mammalian pest control

**DOI:** 10.1101/2023.05.30.542751

**Authors:** Anna C Clark, Alana Alexander, Rey Edison, Kevin Esvelt, Sebastian Kamau, Ludovic Dutoit, Jackson Champer, Samuel E Champer, Philipp W Messer, Neil J Gemmell

## Abstract

1. Fertility-targeted gene drives have been proposed as an ethical genetic approach for managing wild populations of vertebrate pests for public health and conservation benefit.
2. This manuscript introduces a framework to identify and evaluate target gene suitability based on biological gene function, gene expression, and results from mouse knockout models.
3. This framework identified 16 genes essential for male fertility and 12 genes important for female fertility that may be feasible targets for mammalian gene drives and other non-drive genetic pest control technology. Further, a comparative genomics analysis demonstrates the conservation of the identified genes across several globally significant invasive mammals.
4. In addition to providing important considerations for identifying candidate genes, our framework and the genes identified in this study may have utility in developing additional pest control tools such as wildlife contraceptives.

## 1 INTRODUCTION

Invasive vertebrates pose a major global threat to natural ecosystems (Norbury et al., 2013), economies (Latham et al., 2020), and human health (Tait et al., 2017). Pest control can either be achieved through lethal measures or birth control. Traditional physical and chemical pest control approaches typically rely on lethality, often in ways that raise welfare concerns (Littin et al., 2004; Kirk et al., 2020). Lethal control approaches also raise concerns about off-target impacts (Eason et al., 2002, 2011), and efficacy (e.g., resistance to control measures; Anderson et al., 2016; Campbell et al., 2015). Thus, new pest control technology targeting the fertility of a population may gain a higher degree of social acceptance, particularly when combined with genetic engineering techniques (Kirk et al., 2020; Wilkinson & Fitzgerald 2006). This has motivated the development of novel molecular biocontrol approaches, such as meiotic drives that bias the inheritance of particular genes (Burt, 2003). There are several types of meiotic drives, including those that exist in nature (e.g. t-haplotype sex-ratio distorter in mice; Hermann et al., 1999) and others that may be engineered such as homing (Kyrou et al., 2018) and toxin-antidote (Champer et al., 2020) gene drive systems.

Engineered homing gene drives harness guide RNA (gRNA) molecules to direct germline activated endonucleases (e.g., CRISPR-associated Cas9; Pfitzner et al., 2020) to sequences on the wildtype chromosome where they induce a double-strand break (DSB; McFarlane et al., 2018). For population eradication, a homing suppression gene drive can be designed to disrupt genes critical for survival or reproduction. With the drive allele serving as a repair template, homology-directed repair (HDR) allows the drive to be directly inserted into the target gene (Fig. 1A), resulting in both chromosomes and consequent gametes carrying an independent gene drive cassette. Alternatively, disruption through end-joining repair can introduce insertions and deletions at the DSB site to render the target gene non-functional (Fig. 1A), though such alleles often reduce the overall efficiency of the drive. If the transmission rate is sufficiently high, gene drives should spread to high frequency (Fig. 1B), resulting in eventual population collapse if the drive targets genes critical to fertility or fitness (Kyrou et al., 2018).

**Figure 1.**
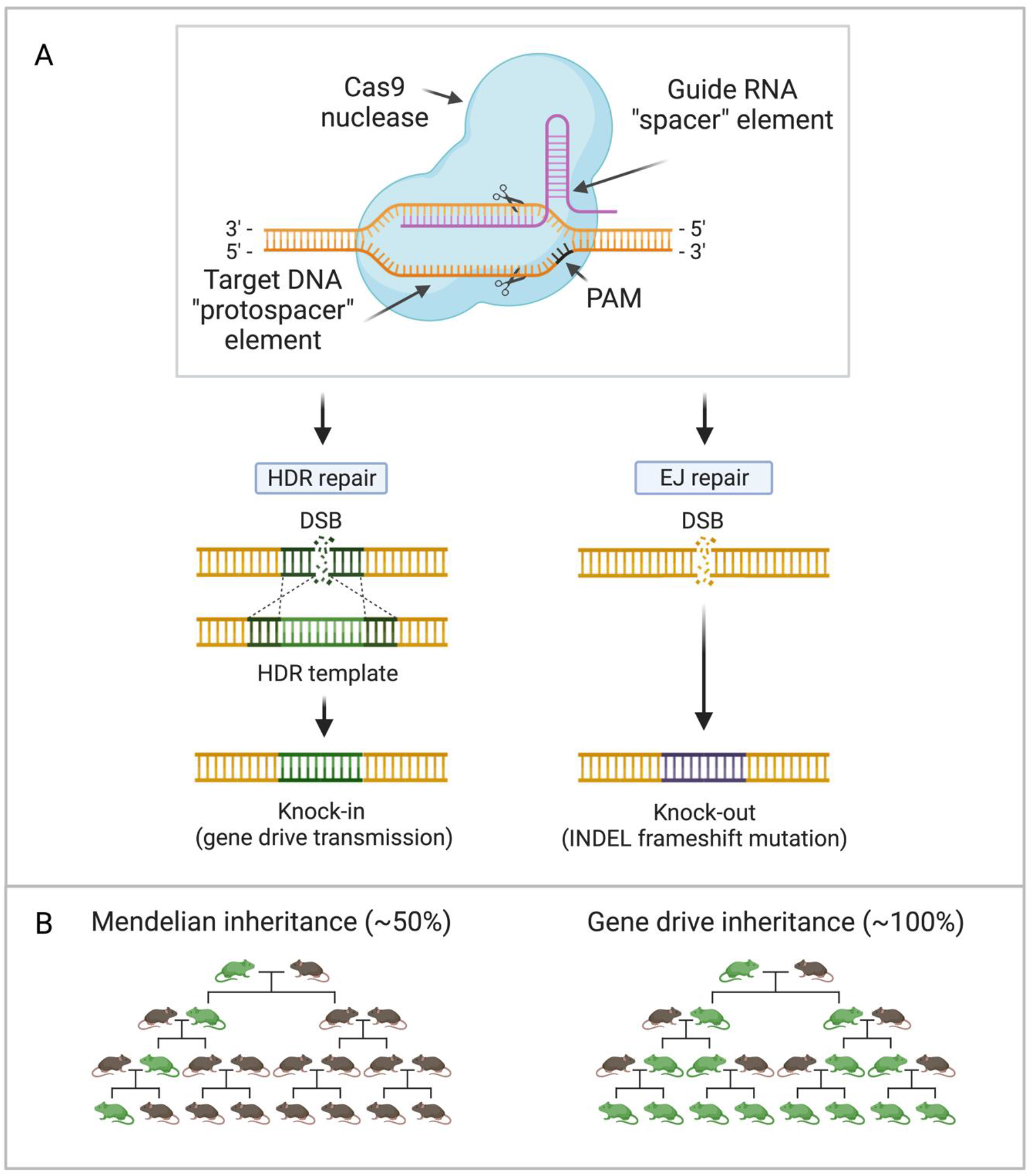
(A) Two mechanisms that homing suppression gene drives use to disrupt genes critical for survival and reproduction: homology-directed repair (HDR; left) and disruption via end joining (EJ; right). In both systems, gRNAs initially target nucleases to a specific genomic locus to induce a double-strand break. If a repair template is provided, HDR leads to insertion of the entire drive construct, including the nuclease, gRNAs, and cargo gene (optional), into the targeted site on the homologous chromosome. Activation of end joining repair, an error prone process, will usually knock-out the gene. EJ repair may thus produce resistance alleles that either conserve or disrupt gene function. Resistance alleles that conserve gene function are problematic as they may prevent the success of future homing drive activity (due to mutations disrupting gRNA target site recognition). (B) Perfect gene drive transmission biases drive allele inheritance in offspring by 100%. Figure from Clark (2023) (CC BY 4.0).

The most important challenges of homing drive systems are the potential evolution of resistance and need for population containment. Resistance to the gene drive can evolve either through standing genetic variation (e.g. mutations in the gRNA target sequences), or drive-induced mutations (e.g. introduction of indels via end joining where HDR was required to transmit the gene drive construct to the homologous chromosome). Function-preserving resistance alleles are particularly concerning in the context of a suppression drive as such alleles can escape cleavage by the drive allele (Champer et al., 2018). Containment is another challenge for homing gene drives as even a small number of drive-carrying individuals are theoretically capable of spreading the drive to an entire population, even if there is a fitness cost to the drive carrier (Esvelt et al., 2014). This means that a drive would likely impact non-target populations, creating significant ecological, social-cultural and political challenges (Taitingfong et al., 2022).

Toxin-antidote systems are a promising method to localise drive spread and reduce the evolution of drive-induced resistance (Champer, Lee, et al., 2020; Champer, Kim, et al., 2020; Champer, Champer, et al., 2021). These drives work by cleaving a gene critical for survival or reproduction (representing the “toxin” portion of the drive), while providing a recoded but functionally equivalent version of the critical target gene (representing the “antidote”), which is linked to the drive construct. The toxin-antidote mechanism thus prevents the persistence of resistance alleles because only individuals that inherit the drive construct can successfully reproduce. Toxin-antidote drives additionally must reach a threshold frequency in the population before they are able to spread (e.g. 2-locus TA gene drives must reach ∼20% frequency; Champer, Champer, et al., 2021). These high rates of initial release could act as a barrier to the drive escaping to non-target populations.

One challenge with toxin-antidote drives is that they can usually only be used for population suppression when used as a tethering system for a paired homing suppression drive (Dhole et al., 2019; Metzloff et al., 2022). One exception to this requires the use of haplolethal target genes, which poses substantial engineering difficulties (non-viability based on resistance allele formation and cell mosaicism caused by CRISPR cleavage during the transformation process (Champer, Yang, et al., 2020)). In either case, toxin-antidote drive versions with the most stringent confinement require relatively high rates of initial release (e.g. >61% initial introduction frequency required by a TADE underdominance suppression drive; Champer, Champer, et al., 2021). Eradication of ship rats from New Zealand using a drive system that requires even just a 5-10% initial inoculation rate would still require the breeding, engineering, and release of tens of thousands of drive carriers, which is an ecologically and economically unrealistic proposition, though it may be possible to release the drive in a limited area and allow it to spread over the landscape.

However, regardless of the type of suppression drive system, there is a research gap in the identification of biologically and ethically suitable candidate genes that could be used to regulate mammalian invasive species. Identifying suitable target fertility genes is also a critical step in investigating and understanding the complex molecular challenges that can arise during gene drive development (e.g., genomic resistance and off-target mutagenesis, and predicting successful *in vivo* application; Rode et al., 2019), as well as to inform parameterisation in predictive gene drive models.

Transgenic expression of sex-determining genes (e.g., *Sry*) has been discussed as a potential method to create a pool of infertile intersex phenotypes, reducing the proportion of fertile females in mammalian populations (Akiyama et al., 2002; Prowse et al., 2017). While promising in theory, experiments have proven challenging and feasibility concerns have been raised (Esvelt & Gemmell, 2017). A more practical approach is to identify and disrupt a functionally important gene region with a specific role in fertility. An ideal target would be a haplosufficient gene essential for fertility in one sex, such that the other sex can transmit the drive further. Targeting a haplosufficient gene essential for fertility in both sexes is not desirable, as such a drive is more likely to prematurely self-extinguish in a large population that is distributed across a heterogenous landscape, particularly if the species dispersal rate and population density is low (Champer, Kim, et al., 2021). Such a drive would instead need to have a reduced genetic load (e.g. targets a gene that only partially reduces fertility), which likely requires higher transmission efficiency to have sufficient power to eradicate the target population (e.g. higher dispersal propensity of drive carriers and high rates of perfect HDR (Fig. 1A)).

Here, we established rigorous criteria involving biological gene function, gene expression, and results from mouse gene knockout (KO) models to identify potential fertility target genes that could be biologically suitable (i.e., those with minimal known off-target effects) for rodent population management using gene drives. These criteria were subsequently used to identify 16 genes indispensable for male fertility and 12 for female fertility that are strong targets for a population suppression drive. We further demonstrated that several of these genes have sequence conservation, which may permit these genes to be investigated and targeted for the development of gene drive systems in other invasive mammalian species. The complexity of gene drive modelling and the expense of laboratory experimentation means that the use of this framework for rigorous pre-screening of potential fertility targets will represent an efficiency multiplier for researchers seeking to develop gene drive technology for the management of invasive species.

## 2 MATERIALS AND METHODS

### 2.1 Gene identification and evaluation

#### 2.1.1 Initial gene identification

We developed and applied a gene evaluation framework to systematically identify candidate fertility genes for mammalian gene drives (Appendix 1). Using the Online Mendelian Inheritance of Man (OMIM) database, we tested and refined keyword searches for genes relating to mammalian gametes or zygote-specific fertility. We used OMIM for initial gene identification because it is a comprehensive resource containing published literature on disease states and loss-of-function mutation causes of mammalian infertility (Hamosh et al., 2005). *Izumo1*, a gene previously identified as specifically essential for male fertility, was utilised as a positive control. The final optimised search yielded a list of 172 genes with the following keyword string:

> (infertil* OR sterility OR sterile OR inviable OR inviability) AND (sperm OR ovum OR egg OR gamete OR zygote) AND (genetic OR gene OR allele) AND (human OR mammal OR mouse OR rat)

We also added a list of 19 genes identified as indispensable for fertility in mice by the Reproductive Genomics Program at the Jackson Laboratory for Mammalian Genetics (hereafter referred to as the Schimenti and Handel gene list; Schimenti and Handel 2018). Gene names, OMIM access number, phenotype, key reference and target sex were recorded. For uniformity, reported gene names and symbols are consistent with nomenclature approved by the HUGO Gene Nomenclature Committee for the gene ortholog in the respective species (Yates et al., 2017). Where there were doubts about the suitability of a gene (e.g., suggestive evidence of functional redundancy or conflicting phenotyping results), genes were excluded.

#### 2.1.2 Inclusion and exclusion criteria

Following the initial gene identification, the gene list was further systematically checked against three criteria defined below in order to identify suitable gene targets for mammalian gene drives. Descriptions and key references for each gene that satisfied our inclusion criteria were further revised and updated in response to new literature.

##### 1. Sex-specific fertility phenotype

Candidate genes must affect gamete and pre-implantation phenotypes to limit instances of individuals carrying moribund embryos and ethical issues surrounding terminating embryonic development and the production of organisms with significant health defects. Evidence of a complete sex-biased fertility phenotype is required to permit the opposite sex to transmit the drive. Summaries of published literature provided by OMIM and additional peer-reviewed literature were reviewed to exclude genes leading to sub-fertility or minor fertility phenotypes in the non-targeted sex.

It is important to note that there are biological similarities between infertility and cancer pathologies, where components of both conditions result from failure to maintain control of somatic and germ cell proliferation (Nagirnaja et al., 2018; Tarín et al., 2015). In cancer, these associated genes are significantly overexpressed, so drives removing function and eliminating gene activity are unlikely to be associated with increased cancer susceptibility. Nevertheless, we excluded genes with evidence of tumour complications in knock-out mice, as this additional source of mortality could have implications on drive performance and ethical acceptability.

##### 2. Tissue-specific expression pattern

Gene expression (assessed via RNAseq data) was required to be restricted or highly overexpressed in specific reproductive tissue (testis or ovary), although minor expression elsewhere was permitted. This criterion reduces the potential for unanticipated phenotypic effects in off-target tissues, which could reduce the fitness of drive-propagating individuals and consequently limit the persistence and spread of the gene drive and may raise ethical concerns. Transcriptome data from the Mouse ENCODE Project via the National Centre for Biotechnology Information (NCBI) online database and Gene Expression Atlas from 23^rd^ September 2018 to 10th October 2018 was used to assess gene expression patterns in different adult tissues in relation to functional pathways, cell type, and developmental stage (Geer et al., 2010; Yue et al., 2014). To construct a quantitative threshold for expression outside the central reproductive tissue, we compared the gene expression results from *Dmrt1* and *Amh*, two genes with strong effects on sexual phenotype (SI Appendix 2; Petryszak et al., 2016; Soumillon et al., 2013). Based on this, genes with non-target tissue expression levels exceeding 5% of the expression levels in target reproductive tissue were excluded to reduce complications of pleiotropic genes.

##### 3. Knock-out mouse model

Raw data from *in vivo* mouse knock-out experiments for each gene of interest were compiled to validate gene expression profiles and support gene function predictions, particularly where it was difficult to isolate and quantify if gene activity in a specific tissue corresponded to an observable phenotype. *In vitro* data was also collected to support the phenotypic observations from *in vivo* experiments. To detect potential sub-fertility phenotypes and intermediate fecundity phenotypes segregating with heterozygotes for each gene, *t*-tests were employed to interrogate differences in litter sizes between genotypes (SI Appendix 3). We investigated this because drive-propagating individuals carrying a single null gene copy are required to be viable with no overt somatic developmental phenotypes to ensure subsequent drive inheritance. The histological impact of the gene knock-outs in specific reproductive tissue (e.g., reduced testis size) was noted. Still, only genes that failed to produce healthy knock-out (KO) offspring were excluded. The absence of potential somatic phenotypes, overt development, abnormal behaviour, and viability between genotypes was confirmed via the literature or through direct correspondence with the original studies’ authors.

#### 2.1.3 Female fertility genes

A further 23 female candidate genes were identified by collaborators using the Mouse Genome Informatics (MGI) between August 2017 and April 2019. Various search terms specifically relating to “fertility defect” were used. Following initial identification, these genes were put through a series of filters, optimising for genes where homozygous mutant female mice displayed near or complete sterility, while heterozygous females and carrier males displayed mostly wild-type fertility. Other exclusion criteria included parturition defects (death of pups during childbirth) and abnormal somatic phenotypes resulting in a significantly reduced fitness of that genotype. When compared to our gene list, many of these female fertility genes had been excluded due to broad somatic expression profiles (i.e., failing our ‘tissue-specific expression pattern’ criteria). To examine the impact of ‘relaxing’ this specific criteria, we instead evaluated these 23 female fertility genes based on female-specific, non-syndromic sterility and the availability of a mouse KO model, ultimately resulting in a list of female target loci (hereafter referred to as the female candidate gene list).

### 2.2 Evolutionary conservation

Invasive vertebrates exist in many terrestrial ecosystems. However, the majority are non-model organisms for which a reference genome has only recently been assembled (stoat and brushtail possum; *unpublished data*, *D. Bond*) or are yet to be sequenced. Where reference sequences are available, conservation within coding regions in well-studied mammalian fertility genes may indicate functional conservation that can be further interrogated once population datasets are available (i.e. identification of variation related to species divergence, and investigation of sites that are near fixed in the population of interest, which may enable population-specific gRNA design for multiple species). Identifying conserved genes would also provide a starting point for empirical experiments, and subsequent tailoring of drive type to the requirements of a particular species. Targeting the same, well characterised gene in multiple species will also be useful to reduce overall resource expenditure for drive development across a number of vertebrate pests. As an island nation with multiple pest species, we use Aotearoa New Zealand (NZ) as an example system to explore the utility of our framework for potentially assisting with gene drive development in the future.

#### 2.2.1 Sequence extraction and processing

Genome assemblies and corresponding gene feature files (GFF) of nine important invasive species in NZ: house mouse (*Mus musculus*), Norway rat (*Rattus norvegicus*), ship rat (*Rattus rattus*), rabbit (*Oryctolagus cuniculus*), western European hedgehog (*Erinaceus europaeus*), domestic cat (*Felis catus*), domestic ferret (*Mustela putorius furo*), stoat (*Mustela erminea*), and common brushtail possum (*Trichosurus vulpecula*), were retrieved from NCBI on 5th November 2020 (SI Appendix 4). Genomic coordinates for each Gnomon-predicted coding sequence (CDS) region within our identified candidate genes were extracted from the GFFs for each species (SI Appendix 5; Souvorov et al., 2010). The extracted coordinates were converted to zero-based BED formatted coordinates using BEDTools (v2.29.2; Quinlan 2014) and used to extract the corresponding nucleotide sequence. As it is optimal for gRNA multiplexing to have multiple adjacent target sites, CDS sequences less than 80 nt were omitted, to allow an optimal minimum of 3 gRNAs of 20 nt each spaced 10 nt apart (CRISPR HDR-based gene drives exhibit optimum drive conversion efficiency with 2-4 gRNAs, and there must be a sufficient number of gRNAs to avoid functional resistance alleles; Champer et al., 2018). Where genes were not present in their respective GFF, a manual search of NCBI for synonymous gene names was conducted to verify the absence of the gene ortholog from the respective target species or to determine the species-specific gene name. Subsequently, the gene feature coordinates for the *Zpbp* ortholog in the brushtail possum were manually added to our dataset.

#### 2.2.2 Multiple sequence alignment

We used LASTZ (1.04.03; Harris 2007) to perform pairwise alignments of coding regions within our candidate genes of interest (Fig. 2; SI Appendix 6). Gap opening was allowed to capture insertions and deletions that may result from species divergence, and the default seed pattern was used. Nucleotide variation was limited to five mismatches per 20 nt during the gapped extension of seed hits to high-scoring pairs (HSP). Back-end filtering of the resulting pairwise alignment blocks set a minimum sequence identity of 75%, consistent with the tolerance of CRISPR-Cas9 gRNAs to no more than five nucleotide mismatches per 20 nt gRNA target sequence (Zheng et al., 2017).

**Figure 2.**
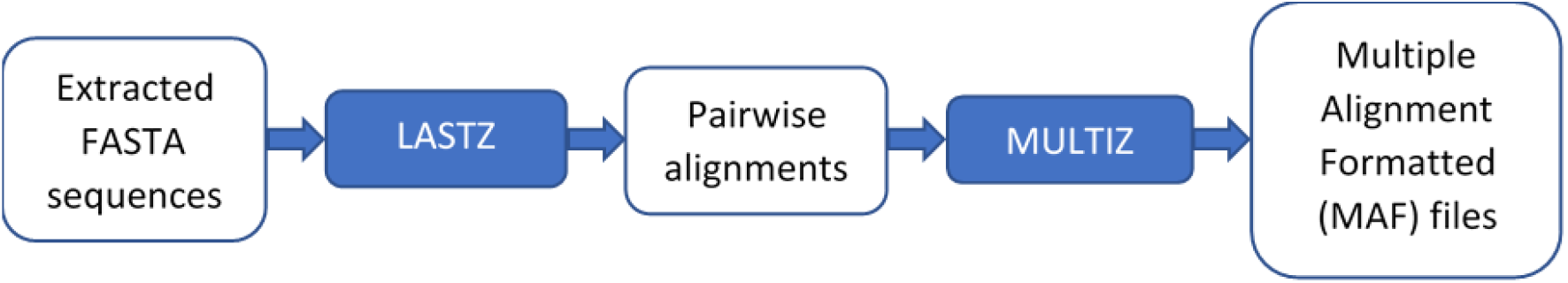
The sequence of steps used to generate and process pairwise alignments, which were subsequently processed into the multiple alignment format. Gene feature files were used to extract coding sequences from whole-genome assemblies. LASTZ (1.04.03) was then used to generate pairwise alignments with at least 75% identity and to limit mismatches to 5 per 20 nt. MULTIZ (v10.6) then filtered the resulting output to eliminate alignments less than 80 nt, and generated the final multiple species local alignments.

The subsequent alignments were then processed by MULTIZ (v10.6; sourced from https://github.com/multiz/multiz.git) to generate a multiple species local alignment of sequence blocks of more than 80 nt (SI Appendix 7). Each conserved sequence block was visualised using tidyverse (v1.3.0; SI Appendix 8; Wickham et al., 2019). Gap openings, including the ends of the alignments, were processed as mismatch penalties, due to the impact of a “missing sequence” on the efficacy of gRNAs designed for these sequence blocks.

## 3 RESULTS

### 3.1 Gene identification

#### 3.1.1 Evaluation and characterisation

Our initial OMIM gene search generated a list of 172 results (SI Appendix 9), to which we added 19 genes from the Schimenti and Handel list to broaden the scope of our investigation (SI Appendix 10; Schimenti & Handel 2018). From the total of 191 results, eight were found to be duplicates (e.g., alternative allelic variants). These entries were merged with their respective genes, generating a list of 183 genes. Application of our filtering criteria identified that 103 genes demonstrated incomplete evidence of a sex-specific phenotype and were subsequently excluded. Sixteen of the remaining 80 genes are specifically critical for male fertility, contributing to several important pre-fertilisation processes, including gametogenesis, gamete recognition, and chromatin remodelling (Table 1; see SI Appendix 10 and 11 for excluded genes). Gene names are consistent with the *Mus musculus* orthologs reported by HGNC. However, variation in nomenclature usage across bodies of literature and databases is evident.

**Table 1.** The 16 genes essential for male fertility identified via our framework. Gene name, OMIM access number, a brief functional description, chromosome number in relation to house mouse (*Mus musculus;* GRCm39: RefSeq assembly accession = GCF_000001635.27) and key functional reference are shown. References: 1. Geyer et al., 2009; 2. Ren & Xia, 2010; 3. Pierre et al., 2012; 4. Miki et al., 2004; 5. Inoue et al., 2005; 6. Zhang et al., 2009; 7. Tokuhiro et al., 2009; 8. Liu et al., 2020, Miyata et al., 2015; 9. Rode et al., 2012; 10. Fujihara et al., 2012; 11. Shao et al., 1999; 12. Zheng et al., 2007; 13. Shang et al., 2017; 14. Spiridonov et al., 2005; 15. Lin et al., 2007.

| Gene | OMIM | Phenotype | Chr | Ref |
| --- | --- | --- | --- | --- |
| *Capza3* | 608722 | Testis-specific actin-capping protein that controls actin polymerisation during spermiogenesis. Expressed in spermatids (Shima et al., 2004). | 6 | 1 |
| *Catsper1* | 606389 | Two of four primary subunits in a sperm-specific Ca<sup>2+</sup> ion channel located in the flagellar, required for the hyperactivation of sperm. Expression is restricted to testes, specifically in developing spermatids, however, expression is not detected in mature sperm (Carlson et al., 2005). | 19 | 2 |
| *Catsper2* | 607249 | “ “ “ “ “ “ | 2 | 2 |
| *Dpy19l2* | 613893 | Spermatid structure between acrosomal and nuclear membranes. Expression is specific to spermatids. | 9 | 3 |
| *Gapdhs* | 610712 | Enzyme localising to the principal piece of the flagellar and involved in energy supply for sperm propulsion. Expression restricted to spermatogenesis. | 7 | 4 |
| *Izumo1* | 609278 | Gamete recognition protein on the sperm cell surface that interacts with the oocyte *Izumo1r* receptor. | 7 | 5 |
| *Meig1* | 614174 | Involved in protein recruitment for sperm flagellar formation. | 2 | 6 |
| *Oaz3* | 605138 | Mammalian antizyme localising to developing spermatids that inhibits polyamine uptake and ornithine decarboxylase activity. Expressed in haploid germ cells. Deficiency results in a separated head and tail of sperm. | 3 | 7 |
| *Ppp3r2* | 613821 | Protein phosphatase regulatory subunit that is expressed throughout sperm development. Deficiency results in inflexible sperm midpiece, sperm motility and morphological defects. | 4 | 8 |
| *Slc26a8* | 608480 | Chloride/sulfate exchange in germ cells and required for sperm motility. Proteins localize in spermatocyte membranes and naïve spermatids. | 17 | 9 |
| *Spaca1* | 612739 | Transmembrane protein localising in the equatorial segment of mature sperm. Expression associated with the inner and outer acrosomal membranes. | 4 | 10 |
| *Spag4* | 603038 | Works with KASH proteins to couple sperm nuclear structures to the cytoskeleton. Expressed in haploid male germ cells. | 2 | 11 |
| *Spem1* | 615116 | Plays a role in detaching cytoplasm from the spermatid nucleus and flagellum neck region required for straightening and stretching of the sperm head and neck. Expressed late in spermatid development. | 11 | 12 |
| *Sun5* | 613942 | Located in the inner membrane of the sperm nuclear envelope and is the most basal protein for flagellar anchoring. First expressed during spermiogenesis (haploid spermatids; Shang et al., 2017). | 2 | 13 |
| *Tssk6* | 610712 | Serine/threonine protein kinase that is required for post-meiotic chromatin remodelling. Expression in spermatids, not spermatocytes (Jha et al., 2017). Another member of this family, *Tssk3*, has been demonstrated to produce a more dramatic phenotype in KO mice (Nayyab et al., 2021). | 8 | 14 |
| *Zpbp* | 608498 | Secondary binding between acrosome-reacted sperm and the zona pellucida. Expressed in spermiogenesis (haploid stages of sperm development). Heterozygous males have an intermediate phenotype (increased levels of abnormal sperm, however, fecundity is comparable to WT sires). | 11 | 15 |

#### 3.1.2 Evaluation of the female fertility genes

The externally contributed list of 23 genes with female-specific phenotypes was evaluated using mouse knock out that demonstrated evidence of a sex-biased sterility phenotype – no tissue-specific expression criteria were applied (see Methods). Eight genes were found to demonstrate either a sub-fertile or potentially lethal phenotype and were subsequently removed from further analysis. Many of the remaining 12 genes (Table 2) have functional roles in maternal effect, gamete recognition, cell signalling, and gene regulatory processes at various stages before embryonic implantation. Two of these genes, *Afp* and *Pgr*, emerged as strong candidates for HDR-based gene drives, despite expression patterns outside the germline (Table 2).

**Table 2.**
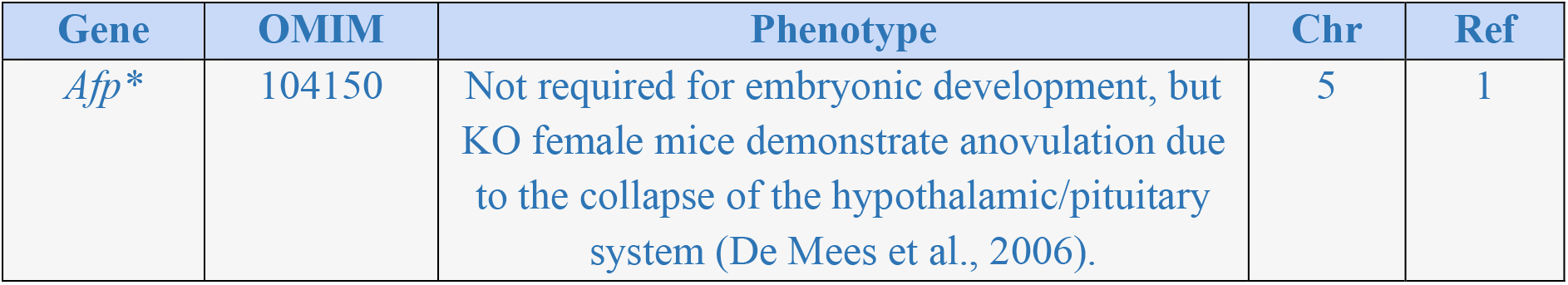
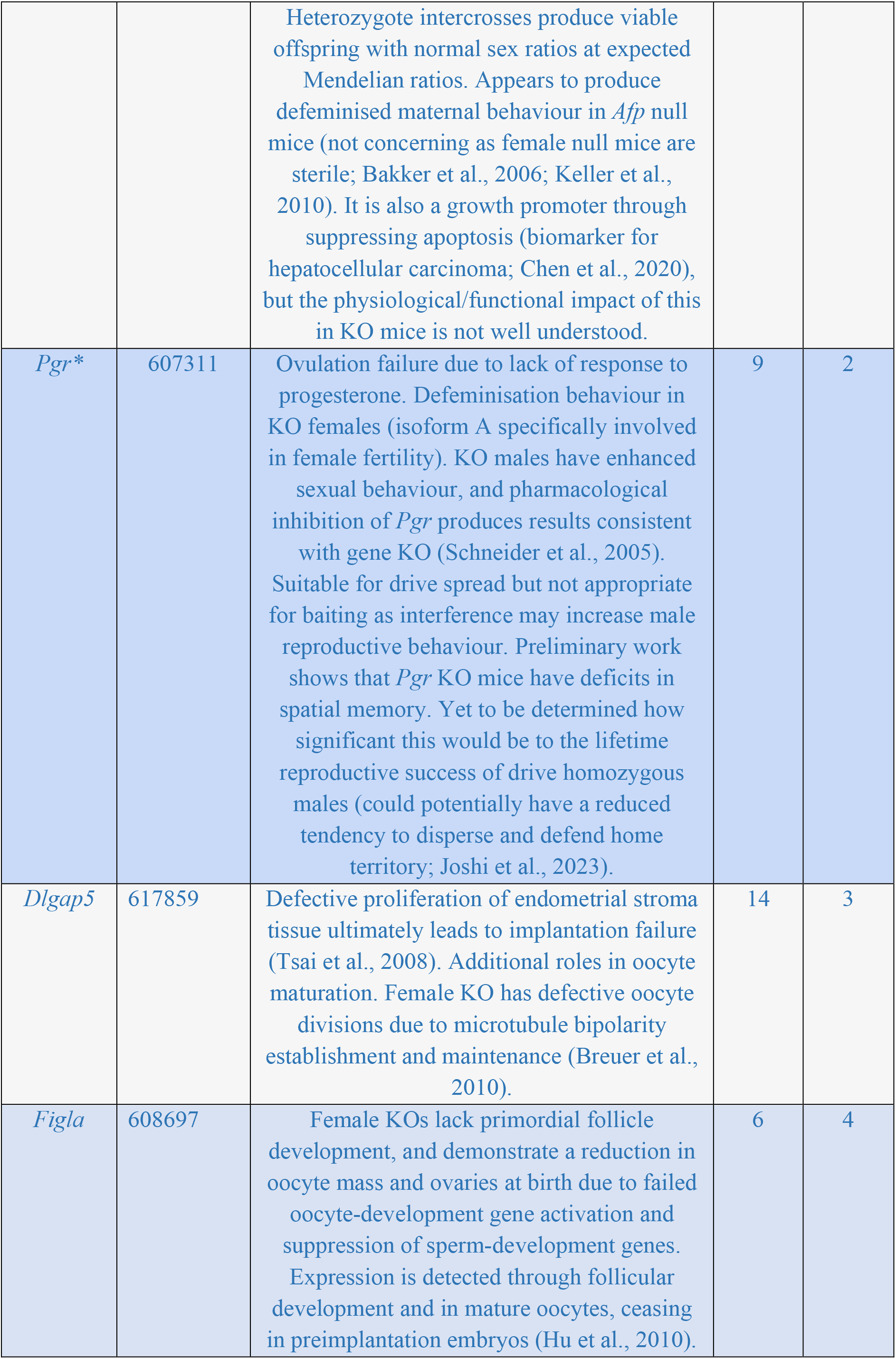
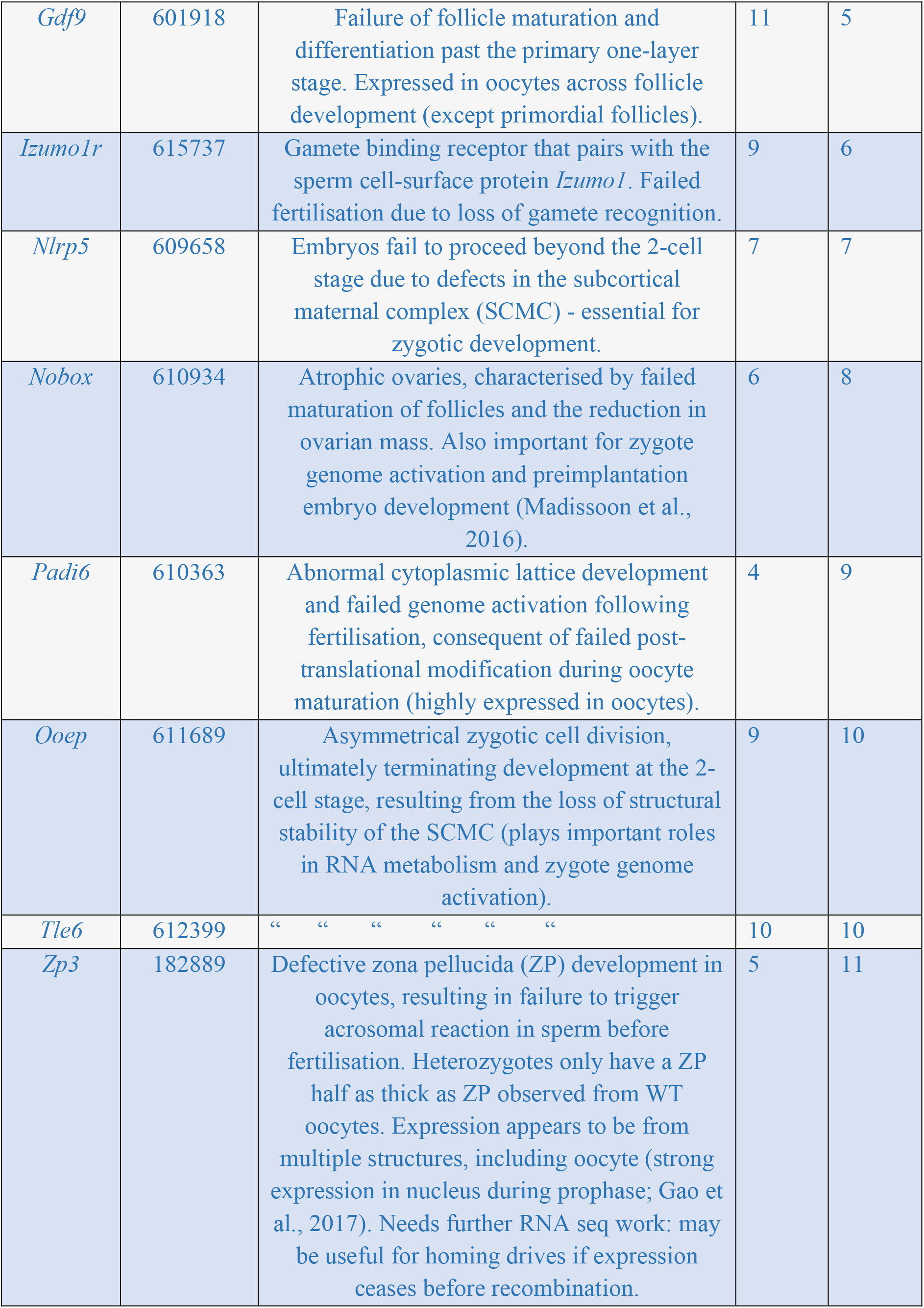
The 12 genes essential for female fertility identified through our evaluation. Asterisks label two genes with somatic expression, which are therefore strong candidates for HDR-based gene drives controlled by a germline promoter. Gene name, OMIM accession number, a brief functional description, chromosome number with relation to house mouse (*Mus musculus*; GRCm39: RefSeq assembly accession = GCF_000001635.27) and key functional reference are shown. References: 1: Gabant et al., 2002; 2: Mulac-Jericevic et al., 2000; 3. Tsai et al., 2008; 4. Soyal et al., 2000; 5. Dong et al., 1996; 6. Bianchi et al., 2014; 7. Tong et al., 2000; 8. Rajkovic et al., 2004; 9. Yurttas et al., 2008; 10. Yu et al., 2014; 11. Liu et al., 2017.

### 3.2 Evolutionary conservation

#### 3.2.1 Overall patterns of sequence conservation

Our multiple species alignments suggest sequence conservation across the nine evaluated mammalian species, including a marsupial representative (brushtail possum; *unpublished data*, *D. Bond*). Sequences of more than 80 nt and at least 75% identity were identified across 10 of 16 evaluated male target genes (*Catsper1*, *Catsper2*, *Gapdhs*, *Meig1*, *Oaz3*, *Slc26a8*, *Spaca1*, *Spag4*, *Tssk6* and *Zpbp*; SI Appendix 12), and 4 of 12 evaluated female candidates (*Dlgap5*, *Pgr*, *Nobox*, and *Zp3*; SI Appendix 13). However, not every species was necessarily represented in each of these conserved blocks (Appendices 14 and 15).

## 4 DISCUSSION

Our systematic gene assessment and subsequent evaluation identified 16 male fertility genes that are critical for mammalian fertility and are strong candidates for genetic population suppression approaches. Using a modified version of our initial evaluation approach (in which we allow minor expression beyond reproductive tissue), we additionally identified 12 female fertility genes. These genes are involved in various reproductive pathways, including structural and gamete recognition, which are processes preceding fertilisation, making these targets a more ethical choice than those with post-fertilisation functions. Many of these genes have previously been suggested as targets for non-hormonal human contraceptives (e.g., *Ppp3r2;* (Castaneda & Matzuk 2015)), and their significance to mammalian fertility has additionally been identified in reviews of the monogenic causes of mammalian fertility (Jiao et al., 2021; Oud et al., 2019). In the process of identifying these genes, we have established a systematic gene identification framework with broad utility, including identifying biological targets for developing species-specific toxins across various taxa, including insects, plants and mammals.

### Male fertility bias and the shortfall of using expression data

During our systematic search and filtering, we observed a complete bias towards genes that are specifically indispensable for male fertility, which highlights that most female fertility factors appear to have phenotypic effects on males, with the reverse being less common due to the extensive and unique set of spermatogenic genes (Matzuk & Lamb 2002). This is consistent with biases presented by the Reproductive Genomics Program (finding a 15:2 ratio of male-specific to female-specific infertility phenotypes; Matzuk & Lamb 2008, Schimenti & Handel 2018) and the Mouse Genome Informatics (finding a 59:21 ratio of male-specific to female-specific infertility phenotypes; Schimenti & Handel 2018). Through our systematic evaluation process, we discovered that most genes specifically indispensable for female fertility demonstrated gene activity in non-target tissue exceeding our tissue-specific expression threshold of 5% (activity of genes essential to female fertility is broad; Dean & Mank 2016). Such genes were subsequently omitted such as to prevent the selection of any genes that may have multiple biological roles and generate multiple pathological phenotypes in allelic-null individuals.

Excluding genes with high expression in somatic tissues also filtered out genes that are somatically expressed but are deposited in the germ cells where they play a significant role in gamete development, maturation and the zygotic activation processes (e.g., maternal and paternal effect genes). Future searches should consider refining the keyword string to probe for genes with isolated expression in granulosa and Sertoli cells as initial drive spread would likely benefit from heterozygotes of both sexes spreading the drive. Alternatively, expression of target fertility genes must precede drive initiated disruption (i.e., meiosis I; Weitzel et al., 2021) so that heterozygous drive carriers remain fertile. Fine-scale spatial characterisation of these expression windows during gamete development remains a major challenge and is an emerging field of new research as technological capabilities (e.g. single cell sequencing) advance.

The substantial male target bias identified through our systematic search is also problematic for suppression gene drive development and application because homing gene drives are most effective at inducing population suppression when targeting female fertility (Burt 2003; Deredec et al., 2008; Prowse et al., 2017). This is because female reproductive success is the major limiting factor in population growth and viability in most species (Galizi et al., 2016; Robertson et al., 2006; Schliekelman et al., 2005; Wedekind 2002). To alleviate the challenges of using gene expression profiles to characterise function for female fertility genes, we utilised fertility and viability data from established mouse knock-out models to verify that gene function predictions are germline-specific. The production of healthy, viable knock-out mice affirms the absence of developmental consequences to the homozygous mutant mice, which implies the absence of any additional phenotypic effects if the gene is coupled to a gene drive. Validating the fitness impact of the candidate gene in both sexes is essential for predicting drive fitness to inform laboratory and computational experiments.

### Functional characterisation

The variations in methods, sample sizes, and the way in which results are reported across the literature are challenging to manage when verifying knockout phenotypes. Where data were available, we found insignificant differences in the litter size between heterozygotes and wildtype individuals of the target sex, validating the absence of a significant intermediate fitness cost in heterozygotes arising from the disruption of one copy of the target gene. However, it should be noted that these results are limited by the experimental environment and genetic background that was tested. Additionally, both female fertility genes identified here as HDR-based gene drive candidates (*Afp* and *Pgr*) appear to play a role in sexual differentiation of the brain early in development and thus have the potential to produce behavioural differences. Further work is necessary in order to characterise how significant this is to the fitness of homozygous null males and confirm the absence of dosage effects in heterozygous gene drive females to understand behavioural limitations of targeting such genes with a drive. Although variations in genetic background and behavioural phenotypes may be relatively small, their effects may be amplified by the dynamics of a gene drive system in large heterogeneous wild populations. Future work should consider using individuals from target populations of interest in representative microcosm environments to control for local genetic and behavioural variation.

### Evolutionary conservation

Several identified fertility genes demonstrate sequence conservation across multiple mammalian lineages, often spanning millions of years of evolutionary divergence. This suggests functional conservation, which may also limit the evolution of functional resistance alleles. These findings separate the genes into two distinct lists – those that are species or lineage-specific, and those that are more broadly conserved and thus may have general utility across a range of target species with species-specific gRNAs. Where resources are limited, targeting research into such general utility genes could substantially reduce the overall research and development expenditure across a cohort of invasive species. It will also ensure confidence that the conserved genic regions have a critical function and therefore are excellent targets.

The functional role of top male fertility genes is heavily biased towards sperm structural development and motility before fertilisation, except *Tssk6*, which regulates the expression of other spermatogenic genes through managing post-meiotic chromatin remodelling in maturing spermatids (Spiridonov et al., 2005). This is likely the result of stronger sexual selection and therefore faster evolutionary rates in sperm-associated genes, which is highlighted in our finding that evolutionarily conserved regulatory genes functioning in fertility are more abundant in the female candidate gene list. Targeting these key regulatory proteins that function in gametogenesis presents a method to create knockout redundancy in the gene drive system.

Additional considerations when screening potential candidate genes include testing gRNA cutting efficiency at such conserved target sites. Further, whole genome assessment is necessary to mitigate the risk of off-target mutagenesis where conserved sequences have been evolutionarily duplicated, giving rise to gene families (a set of genes with a homologous sequence origin but divergent molecular function). Variant assessments of target populations will also enable high-resolution characterisation of target sequences and further facilitate assessments of inter- and intra-species conservation, important for considering drive containment and the evolution of resistance to gene drives.

## 5 CONCLUSION

This stringent systematic analysis has identified a list of sex-specific fertility genes that have the potential as candidate genes for population suppression gene drives in multiple mammal species. The identified candidates have some level of evolutionary conservation and may also have utility for further applications, including developing human and wildlife contraceptives. Additionally, we establish a systematic gene identification framework, including key considerations for candidate gene selection, which may also have utility in other applications, including identifying biological targets for developing species-specific toxins. Further work, including tissue expression profiling, gene knockouts, and population analysis for each target species, will be essential to confirm functional specificity and conservation across mammalian reproductive systems.

## DATA AVAILABILITY

Supplementary information and additional data available from the Figshare Digital Repository: https://doi.org/10.6084/m9.figshare.21905646.v1

## ACKNOWLEDGEMENTS

We thank members of the Gemmell and Messer Labs for helpful discussion, and extend our gratitude to the corresponding authors of the original knock-out mouse experiments for additional clarification of experiment methodology and results. The authors wish to acknowledge the use of New Zealand eScience Infrastructure (NeSI) high performance computing facilities, consulting support and/or training services as part of this research. New Zealand’s national facilities are provided by NeSI and funded jointly by NeSI’s collaborator institutions and through the Ministry of Business, Innovation & Employment’s Research Infrastructure programme. URL https://www.nesi.org.nz. ACC and AA are supported by funding from Predator Free 2050 Ltd. ACC is also supported by funding from Fulbright New Zealand, Ministry for Primary Industries, and Ministry for Pacific Peoples (Toloa Tertiary Scholarship). NJG was supported by funding from Predator Free 2050, Genomics Aotearoa, and the University of Otago. PWM is supported by the National Institutes of Health award R01GM127418.

## CONFLICT OF INTEREST

The authors have no conflicts of interest to declare.

## AUTHORS’ CONTRIBUTIONS

ACC, AA and NJG conceived the ideas and designed methodology; ACC, RE, KE and SK collected the data; ACC, AA and LD developed and conducted analyses; ACC, AA and NJG led the writing of the manuscript; JC, SEC, PWM and LD critically reviewed and edited the manuscript. All authors contributed critically to the drafts and gave final approval for publication.

## Notes

https://doi.org/10.6084/m9.figshare.21905646.v1

### Competing Interest Statement

The authors have declared no competing interest.

## REFERENCES

Akiyama, H., Chaboissier, M.-C., Martin, J. F., Schedl, A., & de Crombrugghe, B. (2002). The transcription factor *Sox9* has essential roles in successive steps of the chondrocyte differentiation pathway and is required for expression of *Sox5* and *Sox6*. Genes & Development, 16(21), 2813–2828. https://doi.org/10.1101/gad.1017802

Anderson, D. P., McMurtrie, P., Edge, K.-A., Baxter, P. W. J., & Byrom, A. E. (2016). Inferential and forward projection modeling to evaluate options for controlling invasive mammals on islands. Ecological Applications, 26(8), 2546–2557.

Bakker, J., De Mees, C., Douhard, Q., Balthazart, J., Gabant, P., Szpirer, J., & Szpirer, C. (2006). Alpha-fetoprotein protects the developing female mouse brain from masculinization and defeminization by estrogens. Nature Neuroscience, 9(2), 220–226. https://doi.org/10.1038/nn1624

Bianchi, E., Doe, B., Goulding, D., Wright, G. J., & Wright, G. J. (2014). *Juno* is the egg *Izumo* receptor and is essential for mammalian fertilization. Nature, 508(7497), 483–487. https://doi.org/10.1038/nature13203

Breuer, M., Kolano, A., Kwon, M., Li, C.-C., Tsai, T.-F., Pellman, D., Brunet, S., & Verlhac, M.-H. (2010). HURP permits MTOC sorting for robust meiotic spindle bipolarity, similar to extra centrosome clustering in cancer cells. Journal of Cell Biology, 191(7), 1251–1260. https://doi.org/10.1083/jcb.201005065

Burt, A. (2003). Site-specific selfish genes as tools for the control and genetic engineering of natural populations. Proceedings of the Royal Society B: Biological Sciences, 270(1518), 921–928. https://doi.org/10.1098/rspb.2002.2319

Campbell, K. J., Beek, J., Eason, C. T., Glen, A. S., Godwin, J., Gould, F., Holmes, N. D., Howald, G. R., Madden, F. M., Ponder, J. B., Threadgill, D. W., Wegmann, A. S., & Baxter, G. S. (2015). The next generation of rodent eradications: Innovative technologies and tools to improve species specificity and increase their feasibility on islands. Biological Conservation, 185, 47–58. http://dx.doi.org/10.1016/j.biocon.2014.10.016

Carlson, A. E., Quill, T. A., Westenbroek, R. E., Schuh, S. M., Hille, B., & Babcock, D. F. (2005). Identical phenotypes of *CatSper1* and *CatSper2* null sperm. The Journal of Biological Chemistry, 280(37), 32238–32244. https://doi.org/10.1074/jbc.M501430200

Castaneda, J., & Matzuk, M. M. (2015). Toward a rapid and reversible male pill. Science, 350(6259), 385–386. https://doi.org/10.1126/science.aad4425

Champer, J., Champer, S. E., Kim, I. K., Clark, A. G., & Messer, P. W. (2021). Design and analysis of CRISPR-based underdominance toxin-antidote gene drives. Evolutionary Applications, 14(4), 1052–1069. https://doi.org/10.1111/EVA.13180

Champer, J., Kim, I. K., Champer, S. E., Clark, A. G., & Messer, P. W. (2020). Performance analysis of novel toxin-antidote CRISPR gene drive systems. BMC Biology 2020 18:1, 18(1), 1–17. https://doi.org/10.1186/S12915-020-0761-2

Champer, J., Kim, I. K., Champer, S. E., Clark, A. G., & Messer, P. W. (2021). Suppression gene drive in continuous space can result in unstable persistence of both drive and wild-type alleles. Molecular Ecology, 30(4), 1086–1101. https://doi.org/10.1111/mec.15788

Champer, J., Lee, E., Yang, E., Liu, C., Clark, A. G., & Messer, P. W. (2020). A toxin-antidote CRISPR gene drive system for regional population modification. Nature Communications, 11(1), 1–10. https://doi.org/10.1038/s41467-020-14960-3

Champer, J., Liu, J., Oh, S. Y., Reeves, R., Luthra, A., Oakes, N., Clark, A. G., & Messer, P. W. (2018). Reducing resistance allele formation in CRISPR gene drive. Proceedings of the National Academy of Sciences, 115(21), 5522–5527. https://doi.org/10.1073/pnas.1720354115

Champer, J., Yang, E., Lee, E., Liu, J., Clark, A. G., & Messer, P. W. (2020). A CRISPR homing gene drive targeting a haplolethal gene removes resistance alleles and successfully spreads through a cage population. Proceedings of the National Academy of Sciences, 117(39), 24377–24383. https://doi.org/10.1073/pnas.2004373117

Chen, T., Dai, X., Dai, J., Ding, C., Zhang, Z., Lin, Z., Hu, J., Lu, M., Wang, Z., Qi, Y., Zhang, L., Pan, R., Zhao, Z., Lu, L., Liao, W., & Lu, X. (2020). AFP promotes HCC progression by suppressing the HuR-mediated Fas/FADD apoptotic pathway. Cell Death & Disease, 11(10), Article 10. https://doi.org/10.1038/s41419-020-03030-7

De Mees, C., Laes, J.-F., Bakker, J., Smitz, J., Hennuy, B., Van Vooren, P., Gabant, P., Szpirer, J., & Szpirer, C. (2006). Alpha-fetoprotein controls female fertility and prenatal development of the gonadotropin-releasing hormone pathway through an antiestrogenic action. Molecular and Cellular Biology, 26(5), 2012–2018. https://doi.org/10.1128/MCB.26.5.2012-2018.2006

Dean, R., & Mank, J. E. (2016). Tissue specificity and sex-specific regulatory variation permit the evolution of sex-biased gene expression. The American Naturalist, 188(3), E74–E84. https://doi.org/10.1086/687526

Deredec, A., Burt, A., & Godfray, H. C. J. (2008). The population genetics of using homing endonuclease genes in vector and pest management. Genetics, 179(4), 2013–2026. https://doi.org/10.1534/genetics.108.089037

Dhole, Sumit, Alun L. Lloyd, and Fred Gould. 2019. “Tethered Homing Gene Drives: A New Design for Spatially Restricted Population Replacement and Suppression.” Evolutionary Applications 12(8):1688–1702. doi: 10.1111/eva.12827.

Dong, J., Albertini, D. F., Nishimori, K., Kumar, T. R., Lu, N., & Matzuk, M. M. (1996). Growth differentiation factor-9 is required during early ovarian folliculogenesis. Nature, 383(6600), 531–535. https://doi.org/10.1038/383531a0

Eason, C. (2002). Sodium monofluoroacetate (1080) risk assessment and risk communication. Toxicology, 181–182, 523–530. https://doi.org/10.1016/S0300-483X(02)00474-2

Eason, C., Miller, A., Ogilvie, S., & Fairweather, A. (2011). An updated review of the toxicology and ecotoxicology of sodium fluoroacetate (1080) in relation to its use as a pest control tool in New Zealand. New Zealand Journal of Ecology, 35, 1–20. https://doi.org/10.2307/24060627

Esvelt, K. M., & Gemmell, N. J. (2017). Conservation demands safe gene drive. PLoS Biology, 15(11), e2003850. https://doi.org/10.1371/journal.pbio.2003850

Esvelt, K. M., Smidler, A. L., Catteruccia, F., & Church, G. M. (2014). Concerning RNA-guided gene drives for the alteration of wild populations. ELife, 3(1), e03401.

Fujihara, Y., Satouh, Y., Inoue, N., Isotani, A., Ikawa, M., & Okabe, M. (2012). *SPACA1*-deficient male mice are infertile with abnormally shaped sperm heads reminiscent of globozoospermia. Development, 139(19), 3583–3589. https://doi.org/10.1242/dev.081778

Gabant, P., Forrester, L., Nichols, J., Van Reeth, T., De Mees, C., Pajack, B., Watt, A., Smitz, J., Alexandre, H., Szpirer, C., & Szpirer, J. (2002). Alpha-fetoprotein, the major fetal serum protein, is not essential for embryonic development but is required for female fertility. Proceedings of the National Academy of Sciences of the United States of America, 99(20), 12865–12870. https://doi.org/10.1073/pnas.202215399

Galizi, R., Hammond, A., Kyrou, K., Taxiarchi, C., Bernardini, F., O’Loughlin, S. M., Papathanos, P.-A., Nolan, T., Windbichler, N., & Crisanti, A. (2016). A CRISPR-Cas9 sex-ratio distortion system for genetic control. Scientific Reports, 6(1), 1–5. https://doi.org/10.1038/srep31139

Gao, L.-L., Zhou, C.-X., Zhang, X.-L., Liu, P., Jin, Z., Zhu, G.-Y., Ma, Y., Li, J., Yang, Z.-X., & Zhang, D. (2017). ZP3 is Required for Germinal Vesicle Breakdown in Mouse Oocyte Meiosis. Scientific Reports, 7(1), Article 1. https://doi.org/10.1038/srep41272

Geer, L. Y., Marchler-Bauer, A., Geer, R. C., Han, L., He, J., He, S., Liu, C., Shi, W., & Bryant, S. H. (2010). The NCBI BioSystems database. Nucleic Acids Research, 38(suppl_1), D492–D496. https://doi.org/10.1093/nar/gkp858

Geyer, C. B., Inselman, A. L., Sunman, J. A., Bornstein, S., Handel, M. A., & Eddy, E. M. (2009). A missense mutation in the *Capza3* gene and disruption of F-actin organization in spermatids of repro32 infertile male mice. Developmental Biology, 330(1), 142–152. https://doi.org/10.1016/j.ydbio.2009.03.020

Hamosh, A., Scott, A. F., Amberger, J. S., Bocchini, C. A., & McKusick, V. A. (2005). Online Mendelian Inheritance in Man (OMIM), a knowledgebase of human genes and genetic disorders. Nucleic Acids Research, 33(suppl_1), D514–D517. https://doi.org/10.1093/nar/gki033

Harris, R. S. (2007). Improved Pairwise Alignment Of Genomic DNA. Pennsylvania State University.

Hu, W., Gauthier, L., Baibakov, B., Jimenez-Movilla, M., & Dean, J. (2010). FIGLA, a Basic Helix-Loop-Helix Transcription Factor, Balances Sexually Dimorphic Gene Expression in Postnatal Oocytes. Molecular and Cellular Biology, 30(14), 3661–3671. https://doi.org/10.1128/MCB.00201-10

Inoue, N., Ikawa, M., Isotani, A., & Okabe, M. (2005). The immunoglobulin superfamily protein *Izumo* is required for sperm to fuse with eggs. Nature, 434(7030), 234–238. https://doi.org/10.1038/nature03362

Jha, K. N., Tripurani, S. K., & Johnson, G. R. (2017). *TSSK6* is required for γH2AX formation and the histone-to-protamine transition during spermiogenesis. Journal of Cell Science, 130(10), 1835–1844. https://doi.org/10.1242/jcs.202721

Jiao, S.-Y., Yang, Y.-H., & Chen, S.-R. (2021). Molecular genetics of infertility: Loss-of-function mutations in humans and corresponding knockout/mutated mice. Human Reproduction Update, 27(1), 154–189. https://doi.org/10.1093/humupd/dmaa034

Joshi, S., Williams, C. L., & Kapur, J. (2023). Limbic progesterone receptors regulate spatial memory. Scientific Reports, 13(1), 2164. https://doi.org/10.1038/s41598-023-29100-2

Keller, M., Pawluski, J. L., Brock, O., Douhard, Q., & Bakker, J. (2010). The α-fetoprotein knock-out mouse model suggests that parental behavior is sexually differentiated under the influence of prenatal estradiol. Hormones and Behavior, 57(4–5), 434–440. https://doi.org/10.1016/j.yhbeh.2010.01.013

Kirk, N. A., Kannemeyer, R., Greenaway, A., Macdonald, E., Stronge, D., & Whenua, A. A. M. (2020). Understanding attitudes on new technologies to manage invasive species. Pacific Conservation Biology, 26(1), 35–44. https://doi.org/10.1071/PC18080

Kyrou, K., Hammond, A. M., Galizi, R., Kranjc, N., Burt, A., Beaghton, A. K., Nolan, T., & Crisanti, A. (2018). A CRISPR–Cas9 gene drive targeting doublesex causes complete population suppression in caged Anopheles gambiae mosquitoes. Nature Biotechnology 2018 36:11, 36(11), 1062–1066. https://doi.org/10.1038/nbt.4245

Latham, A. D. M., Latham, M. C., Norbury, G. L., Forsyth, D. M., & Warburton, B. (2020). A review of the damage caused by invasive wild mammalian herbivores to primary production in New Zealand. New Zealand Journal of Zoology, 47(1), 20–52. https://doi.org/10.1080/03014223.2019.1689147

Lin, Y.-N., Roy, A., Yan, W., Burns, K. H., & Matzuk, M. M. (2007). Loss of zona pellucida binding proteins in the acrosomal matrix disrupts acrosome biogenesis and sperm morphogenesis. Molecular and Cellular Biology, 27(19), 6794–6805. https://doi.org/10.1128/MCB.01029-07

Littin, K., Mellor, D., Warburton, B., & Eason, C. (2004). Animal welfare and ethical issues relevant to the humane control of vertebrate pests. New Zealand Veterinary Journal, 52(1), 1–10. https://doi.org/10.1080/00480169.2004.36384

Liu, W., Li, K., Bai, D., Yin, J., Tang, Y., Chi, F., Zhang, L., Wang, Y., Pan, J., Liang, S., Guo, Y., Ruan, J., Kou, X., Zhao, Y., Wang, H., Chen, J., Teng, X., & Gao, S. (2017). Dosage effects of *ZP2* and *ZP3* heterozygous mutations cause human infertility. Human Genetics, 136(8), 975–985. https://doi.org/10.1007/s00439-017-1822-7

Liu, Y., Zhang, C., Wang, S., Hu, Y., Jing, J., Ye, L., Jing, R., & Ding, Z. (2020). Dependence of sperm structural and functional integrity on testicular calcineurin isoform PPP3R2 expression. Journal of Molecular Cell Biology, 12(7), 515–529. https://doi.org/10.1093/jmcb/mjz115

Madissoon, E., Jouhilahti, E.-M., Vesterlund, L., Töhönen, V., Krjutškov, K., Petropoulos, S., Einarsdottir, E., Linnarsson, S., Lanner, F., Månsson, R., Hovatta, O., Bürglin, T. R., Katayama, S., & Kere, J. (2016). Characterization and target genes of nine human PRD-like homeobox domain genes expressed exclusively in early embryos. Scientific Reports, 6(1), Article 1. https://doi.org/10.1038/srep28995

Matzuk, M. M., & Lamb, D. J. (2002). Genetic dissection of mammalian fertility pathways. Nature Medicine, 8(10), S33–S40. https://doi.org/10.1038/nm-fertilityS41

Matzuk, M. M., & Lamb, D. J. (2008). The biology of infertility: Research advances and clinical challenges. Nature Medicine, 14(11), 1197–1213. https://doi.org/10.1038/nm.f.1895

McFarlane, G. R., Whitelaw, C. B. A., & Lillico, S. G. (2018). CRISPR-based gene drives for pest control. Trends in Biotechnology, 36(2), 130–133. https://doi.org/10.1016/j.tibtech.2017.10.001

Metzloff, Matthew, Emily Yang, Sumit Dhole, Andrew G. Clark, Philipp W. Messer, and Jackson Champer. 2022. “Experimental Demonstration of Tethered Gene Drive Systems for Confined Population Modification or Suppression.” BMC Biology 20(1):119. doi: 10.1186/s12915-022-01292-5.

Miki, K., Qu, W., Goulding, E. H., Willis, W. D., Bunch, D. O., Strader, L. F., Perreault, S. D., Eddy, E. M., & O’Brien, D. A. (2004). Glyceraldehyde 3-phosphate dehydrogenase-S, a sperm-specific glycolytic enzyme, is required for sperm motility and male fertility. Proceedings of the National Academy of Sciences of the United States of America, 101(47), 16501 LP – 16506. https://doi.org/10.1073/pnas.0407708101

Miyata, H., Satouh, Y., Mashiko, D., Muto, M., Nozawa, K., Shiba, K., Fujihara, Y., Isotani, A., Inaba, K., & Ikawa, M. (2015). Sperm calcineurin inhibition prevents mouse fertility with implications for male contraceptive. Science, 350(6259), 442–445. https://doi.org/10.1126/science.aad0836

Mulac-Jericevic, B., Mullinax, R. A., DeMayo, F. J., Lydon, J. P., & Conneely, O. M. (2000). Subgroup of reproductive functions of progesterone mediated by progesterone receptor-B isoform. Science, 289(5485), 1751–1754. https://doi.org/10.1126/science.289.5485.1751

Nagirnaja, L., Aston, K. I., & Conrad, D. F. (2018). Genetic intersection of male infertility and cancer. Fertility and Sterility, 109(1), 20–26. https://doi.org/10.1016/j.fertnstert.2017.10.028

Nayyab, S., Gervasi, M. G., Tourzani, D. A., Caraballo, D. A., Jha, K. N., Teves, M. E., Cui, W., Georg, G. I., Visconti, P. E., & Salicioni, A. M. (2021). TSSK3, a novel target for male contraception, is required for spermiogenesis. Molecular Reproduction and Development, 88(11), 718–730. https://doi.org/10.1002/mrd.23539

Norbury, G., Byrom, A., Pech, R., Smith, J., Clarke, D., Anderson, D., & Forrester, G. (2013). Invasive mammals and habitat modification interact to generate unforeseen outcomes for indigenous fauna. Ecological Applications, 23(7), 1707–1721. https://doi.org/10.1890/12-1958.1

Oud, M. S., Volozonoka, L., Smits, R. M., Vissers, L. E. L. M., Ramos, L., & Veltman, J. A. (2019). A systematic review and standardized clinical validity assessment of male infertility genes. Human Reproduction, 34(5), 932–941. https://doi.org/10.1093/humrep/dez022

Petryszak, R., Keays, M., Tang, Y. A., Fonseca, N. A., Barrera, E., Burdett, T., Füllgrabe, A., Fuentes, A. M.-P., Jupp, S., Koskinen, S., Mannion, O., Huerta, L., Megy, K., Snow, C., Williams, E., Barzine, M., Hastings, E., Weisser, H., Wright, J., … Brazma, A. (2016). Expression Atlas update—An integrated database of gene and protein expression in humans, animals and plants. Nucleic Acids Research, 44(D1), D746–D752. https://doi.org/10.1093/nar/gkv1045

Pfitzner, C., White, M. A., Piltz, S. G., Scherer, M., Adikusuma, F., Hughes, J. N., & Thomas, P. Q. (2020). Progress Toward Zygotic and Germline Gene Drives in Mice. The CRISPR Journal, 3(5), 388–397. https://doi.org/10.1089/crispr.2020.0050

Pierre, V., Martinez, G., Coutton, C., Delaroche, J., Yassine, S., Novella, C., Pernet-Gallay, K., Hennebicq, S., Ray, P. F., & Arnoult, C. (2012). Absence of *Dpy19l2*, a new inner nuclear membrane protein, causes globozoospermia in mice by preventing the anchoring of the acrosome to the nucleus. Development, 139(16), 2955–2965. https://doi.org/10.1242/dev.077982

Prowse, T. A. A., Cassey, P., Ross, J. V., Pfitzner, C., Wittmann, T. A., & Thomas, P. (2017). Dodging silver bullets: Good CRISPR gene-drive design is critical for eradicating exotic vertebrates. Proceedings of the Royal Society B: Biological Sciences, 248(1860), 20170799.

Quinlan, A. R. (2014). BEDTools: The Swiss-army tool for genome feature analysis. Current Protocols in Bioinformatics, 47(1), 11–12. https://doi.org/10.1002/0471250953.bi1112s47

Rajkovic, A., Pangas, S. A., Ballow, D., Suzumori, N., & Matzuk, M. M. (2004). NOBOX deficiency disrupts early folliculogenesis and oocyte-specific gene expression. Science, 305(5687), 1157–1159. https://doi.org/10.1126/science.1099755

Ren, D., & Xia, J. (2010). Calcium signaling through CatSper channels in mammalian fertilization. Physiology, 25(3), 165–175. https://doi.org/10.1152/physiol.00049.2009

Robertson, B. C., Elliott, G. P., Eason, D. K., Clout, M. N., & Gemmell, N. J. (2006). Sex allocation theory aids species conservation. Biology Letters, 2(2), 229–231. https://doi.org/10.1098/rsbl.2005.0430

Rode, B., Dirami, T., Bakouh, N., Rizk-Rabin, M., Norez, C., Lhuillier, P., Lorès, P., Jollivet, M., Melin, P., Zvetkova, I., Bienvenu, T., Becq, F., Planelles, G., Edelman, A., Gacon, G., & Touré, A. (2012). The testis anion transporter TAT1 (SLC26A8) physically and functionally interacts with the cystic fibrosis transmembrane conductance regulator channel: A potential role during sperm capacitation. Human Molecular Genetics, 21(6), 1287–1298. https://doi.org/10.1093/hmg/ddr558

Rode, N. O., Estoup, A., Bourguet, D., Courtier-Orgogozo, V., & Débarre, F. (2019). Population management using gene drive: Molecular design, models of spread dynamics and assessment of ecological risks. Conservation Genetics, 20(4), 671–690. https://doi.org/10.1007/s10592-019-01165-5

Schimenti, J. C., & Handel, M. A. (2018). Unpackaging the genetics of mammalian fertility: Strategies to identify the “reproductive genome.” Biology of Reproduction, 99(6), 1119– 1128. https://doi.org/10.1093/biolre/ioy133

Schliekelman, P., Ellner, S., & Gould, F. (2005). Pest control by genetic manipulation of sex ratio. Journal of Economic Entomology, 98(1), 18–34. https://doi.org/10.1093/jee/98.1.18

Schneider, J. S., Burgess, C., Sleiter, N. C., DonCarlos, L. L., Lydon, J. P., O’Malley, B., & Levine, J. E. (2005). Enhanced sexual behaviors and androgen receptor immunoreactivity in the male progesterone receptor knockout mouse. Endocrinology, 146(10), 4340–4348. https://doi.org/10.1210/en.2005-0490

Shang, Y., Zhu, F., Wang, L., Ouyang, Y.-C., Dong, M.-Z., Liu, C., Zhao, H., Cui, X., Ma, D., Zhang, Z., Yang, X., Guo, Y., Liu, F., Yuan, L., Gao, F., Guo, X., Sun, Q.-Y., Cao, Y., & Li, W. (2017). Essential role for SUN5 in anchoring sperm head to the tail. ELife, 6. https://doi.org/10.7554/eLife.28199

Shao, X., Tarnasky, H. A., Lee, J. P., Oko, R., & van der Hoorn, F. A. (1999). *Spag4*, a novel sperm protein, binds outer dense-fiber protein *Odf1* and localizes to microtubules of manchette and axoneme. Developmental Biology, 211(1), 109–123. https://doi.org/10.1006/dbio.1999.9297

Shima, J. E., McLean, D. J., McCarrey, J. R., & Griswold, M. D. (2004). The murine testicular transcriptome: Characterizing gene expression in the testis during the progression of spermatogenesis. Biology of Reproduction, 71(1), 319–330. https://doi.org/10.1095/biolreprod.103.026880

Soumillon, M., Necsulea, A., Weier, M., Brawand, D., Zhang, X., Gu, H., Barthès, P., Kokkinaki, M., Nef, S., Gnirke, A., Dym, M., de Massy, B., Mikkelsen, T. S., & Kaessmann, H. (2013). Cellular source and mechanisms of high transcriptome complexity in the mammalian testis. Cell Reports, 3(6), 2179–2190. https://doi.org/10.1016/J.CELREP.2013.05.031

Souvorov, A., Kapustin, Y., Kiryutin, B., Chetvernin, V., Tatusova, T., & Lipman, D. (2010). Gnomon-NCBI eukaryotic gene prediction tool. National Center for Biotechnology Information, 1–24.

Soyal, S. M., Amleh, A., & Dean, J. (2000). FIGalpha, a germ cell-specific transcription factor required for ovarian follicle formation. Development, 127(21), 4645 LP – 4654.

Spiridonov, N. A., Wong, L., Zerfas, P. M., Starost, M. F., Pack, S. D., Paweletz, C. P., & Johnson, G. R. (2005). Identification and characterization of SSTK, a serine/threonine protein kinase essential for male fertility. Molecular and Cellular Biology, 25(10), 4250– 4261. https://doi.org/10.1128/MCB.25.10.4250-4261.2005

Tait, P., Saunders, C., Nugent, G., & Rutherford, P. (2017). Valuing conservation benefits of disease control in wildlife: A choice experiment approach to bovine tuberculosis management in New Zealand’s native forests. Journal of Environmental Management, 189, 142–149. https://doi.org/10.1016/j.jenvman.2016.12.045

Taitingfong, R. I., Triplett, C., Vásquez, V. N., Rajagopalan, R. M., Raban, R., Roberts, A., Terradas, G., Baumgartner, B., Emerson, C., Gould, F., Okumu, F., Schairer, C. E., Bossin, H. C., Buchman, L., Campbell, K. J., Clark, A., Delborne, J., Esvelt, K., Fisher, J., … Bloss, C. S. (2022). Exploring the value of a global gene drive project registry. Nature Biotechnology, 1–5. https://doi.org/10.1038/s41587-022-01591-w

Tarín, J. J., García-Pérez, M. A., Hamatani, T., & Cano, A. (2015). Infertility etiologies are genetically and clinically linked with other diseases in single meta-diseases. Reproductive Biology and Endocrinology, 13(1), 1–11. https://doi.org/10.1186/s12958-015-0029-9

Tokuhiro, K., Isotani, A., Yokota, S., Yano, Y., Oshio, S., Hirose, M., Wada, M., Fujita, K., Ogawa, Y., Okabe, M., Nishimune, Y., & Tanaka, H. (2009). OAZ-t/OAZ3 Is Essential for Rigid Connection of Sperm Tails to Heads in Mouse. PLoS Genetics, 5(11), e1000712. https://doi.org/10.1371/journal.pgen.1000712

Tong, Z. B., Gold, L., Pfeifer, K. E., Dorward, H., Lee, E., Bondy, C. A., Dean, J., & Nelson, L. M. (2000). *Mater*, a maternal effect gene required for early embryonic development in mice. Nature Genetics, 26(3), 267–268. https://doi.org/10.1038/81547

Tsai, C. Y., Chou, C. K., Yang, C. W., Lai, Y. C., Liang, C. C., Chen, C. M., & Tsai, T. F. (2008). Hurp deficiency in mice leads to female infertility caused by an implantation defect. Journal of Biological Chemistry, 283(39), 26302–26306. https://doi.org/10.1074/jbc.C800117200

Wedekind, C. (2002). Manipulating sex ratios for conservation: Short-term risks and long-term benefits. Animal Conservation, 5(1), 13–20. https://doi.org/10.1017/S1367943002001026

Weitzel, A. J., Grunwald, H. A., Weber, C., Levina, R., Gantz, V. M., Hedrick, S. M., Bier, E., & Cooper, K. L. (2021). Meiotic Cas9 expression mediates gene conversion in the male and female mouse germline. PLoS Biology, 19(12), e3001478. https://doi.org/10.1371/journal.pbio.3001478

Wickham, H., Averick, M., Bryan, J., Chang, W., McGowan, L., François, R., Grolemund, G., Hayes, A., Henry, L., Hester, J., Kuhn, M., Pedersen, T., Miller, E., Bache, S., Müller, K., Ooms, J., Robinson, D., Seidel, D., Spinu, V., … Yutani, H. (2019). Welcome to the Tidyverse. Journal of Open Source Software, 4(43), 1686. https://doi.org/10.21105/joss.01686

Wilkinson, R., & Fitzgerald, G. (2006). Public attitudes toward possum fertility control and genetic engineering in New Zealand (p. 52). Landcare Research New Zealand Limited.

Yates, B., Braschi, B., Gray, K. A., Seal, R. L., Tweedie, S., & Bruford, E. A. (2017). Genenames.org: The HGNC and VGNC resources in 2017. Nucleic Acids Research, 45(D1), D619–D625. https://doi.org/10.1093/nar/gkw1033

Yu, X. J., Yi, Z., Gao, Z., Qin, D., Zhai, Y., Chen, X., Ou-Yang, Y., Wang, Z. B., Zheng, P., Zhu, M. S., Wang, H., Sun, Q. Y., Dean, J., & Li, L. (2014). The subcortical maternal complex controls symmetric division of mouse zygotes by regulating F-actin dynamics. Nature Communications, 5(1), 4887. https://doi.org/10.1038/ncomms5887

Yue, F., Cheng, Y., Breschi, A., Vierstra, J., Wu, W., Ryba, T., Sandstrom, R., Ma, Z., Davis, C., Pope, B. D., Shen, Y., Pervouchine, D. D., Djebali, S., Thurman, R. E., Kaul, R., Rynes, E., Kirilusha, A., Marinov, G. K., Williams, B. A., … Consortium, T. M. E. (2014). A comparative encyclopedia of DNA elements in the mouse genome. Nature, 515(7527), 355–364. https://doi.org/10.1038/nature13992

Yurttas, P., Vitale, A. M., Fitzhenry, R. J., Cohen-Gould, L., Wu, W., Gossen, J. A., & Coonrod, S. A. (2008). Role for PADI6 and the cytoplasmic lattices in ribosomal storage in oocytes and translational control in the early mouse embryo. Development, 135(15), 2627–2636. https://doi.org/10.1242/dev.016329

Zhang, Z., Shen, X., Gude, D. R., Wilkinson, B. M., Justice, M. J., Flickinger, C. J., Herr, J. C., Eddy, E. M., & Strauss, J. F. (2009). MEIG1 is essential for spermiogenesis in mice. Proceedings of the National Academy of Sciences, 106(40), 17055–17060. https://doi.org/10.1073/pnas.0906414106

Zheng, H., Stratton, C. J., Morozumi, K., Jin, J., Yanagimachi, R., & Yan, W. (2007). Lack of *Spem1* causes aberrant cytoplasm removal, sperm deformation, and male infertility. Proceedings of the National Academy of Sciences, 104(16), 6852–6857.

Zheng, T., Hou, Y., Zhang, P., Zhang, Z., Xu, Y., Zhang, L., Niu, L., Yang, Y., Liang, D., Yi, F., Peng, W., Feng, W., Yang, Y., Chen, J., Zhu, Y. Y., Zhang, L. H., & Du, Q. (2017). Profiling single-guide RNA specificity reveals a mismatch sensitive core sequence. Scientific Reports, 7(1), 1–8. https://doi.org/10.1038/srep40638

